# Assessing Lanthanide-Dependent Methanol Dehydrogenase Activity: The Assay Matters

**DOI:** 10.1101/2023.10.26.563981

**Authors:** Manh Tri Phi, Helena Singer, Felix Zäh, Christoph Haisch, Sabine Schneider, Huub J.M. Op den Camp, Lena J. Daumann

## Abstract

Artificial dye-coupled assays have been widely adopted as a rapid and convenient method to assess the activity of methanol dehydrogenases (MDH). Lanthanide(Ln)-dependent XoxF-MDHs from methanotrophs and methylotrophs are able to incorporate different lanthanides (Lns) in their active site. The artificial dye-coupled assay showed that the earlier Lns exhibit a higher enzyme activity than the late Lns. Although this assay is widely used, there are limitations. It is not unusual that a pH of 9 is required and that activators like ammonium have to be added to the assay mixture which do not reflect the conditions inside the cell. Moreover, different Ln-MDH variants are not obtained by the direct isolation from the cells grown with the respective Ln, but by metal titration of the Ln in a partial-apo-MDH or by incubation of an apo-MDH with the Ln. Herein, we report the cultivation of Ln-dependent methanotroph *Methylacidiphilum fumariolicum* SolV with nine different Lns, the isolation of the respective MDH and the assessment of the enzyme activity using the artificial dye-coupled assay. We compare these results with an adapted protein-coupled assay using the physiological partner and electron acceptor cytochrome *c*_*GJ*_ (cyt *c*_*GJ*_) instead of artificial dyes. We demonstrate that, depending on the assay, two distinct trends are observed among the Ln series. The specific activity of La-, Ce- and Pr-MDH, as measured by the protein-coupled assay, exceeds the specific enzyme activity measured by the dye-coupled assay. This suggests that the early Lns also have a positive effect on the interaction between XoxF-MDH and its cyt *c*_*GJ*_ thereby increasing functional efficiency.

## Introduction

In recent years, lanthanides (Lns, La–Lu) have firmly been established as biological relevant. Lanthanide(Ln)-dependent or -utilizing bacteria have been found in a variety of ecosystems, e.g. phyllosphere, pond sediment, (coastal) marine environment, shale rock, rice rhizosphere and geothermal fields.^1-7^ Most of these bacteria are either methylotrophs or methanotrophs and use small C_1_-molecules like methane, methanol, halogenated methanes, methylated amines and methylated sulfur species as their energy source.^8^ Methanotrophs are able to convert methane to carbon dioxide and play a significant role in the global carbon cycle.^3^ In the first step, methane is oxidized to methanol by particulate methane monooxygenase or soluble methane monooxygenase and subsequently oxidized to formaldehyde by methanol dehydrogenase (MDH).^9-11^ There are two variants of this MDH: Ca-containing MxaFI-MDH and Ln-containing XoxF-MDH. All methanotrophs that possess the MxaFI-MDH variant also have the XoxF-MDH.^8, 12, 13^ There are also reports of methano- and methylotrophs that exclusively possess the XoxF-MDH, highlighting the widespread prevalence of Ln-utilizing microorganisms.^7, 14, 15^ If both types of MDH are present, MxaFI-MDH is expressed in the absence of any Ln. However, even the presence of nanomolar amounts of Ln is sufficient to initiate a transcriptional response, the “lanthanide-switch”, favouring the expression of the XoxF-MDH variant even when the concentration of Ca is 100-fold higher.^16-19^ In addition to the Ln ion, the active site contains pyrroloquinoline quinone (PQQ) as the second essential cofactor for XoxF-MDH (Fig. 1A).^20^

**Figure 1.**
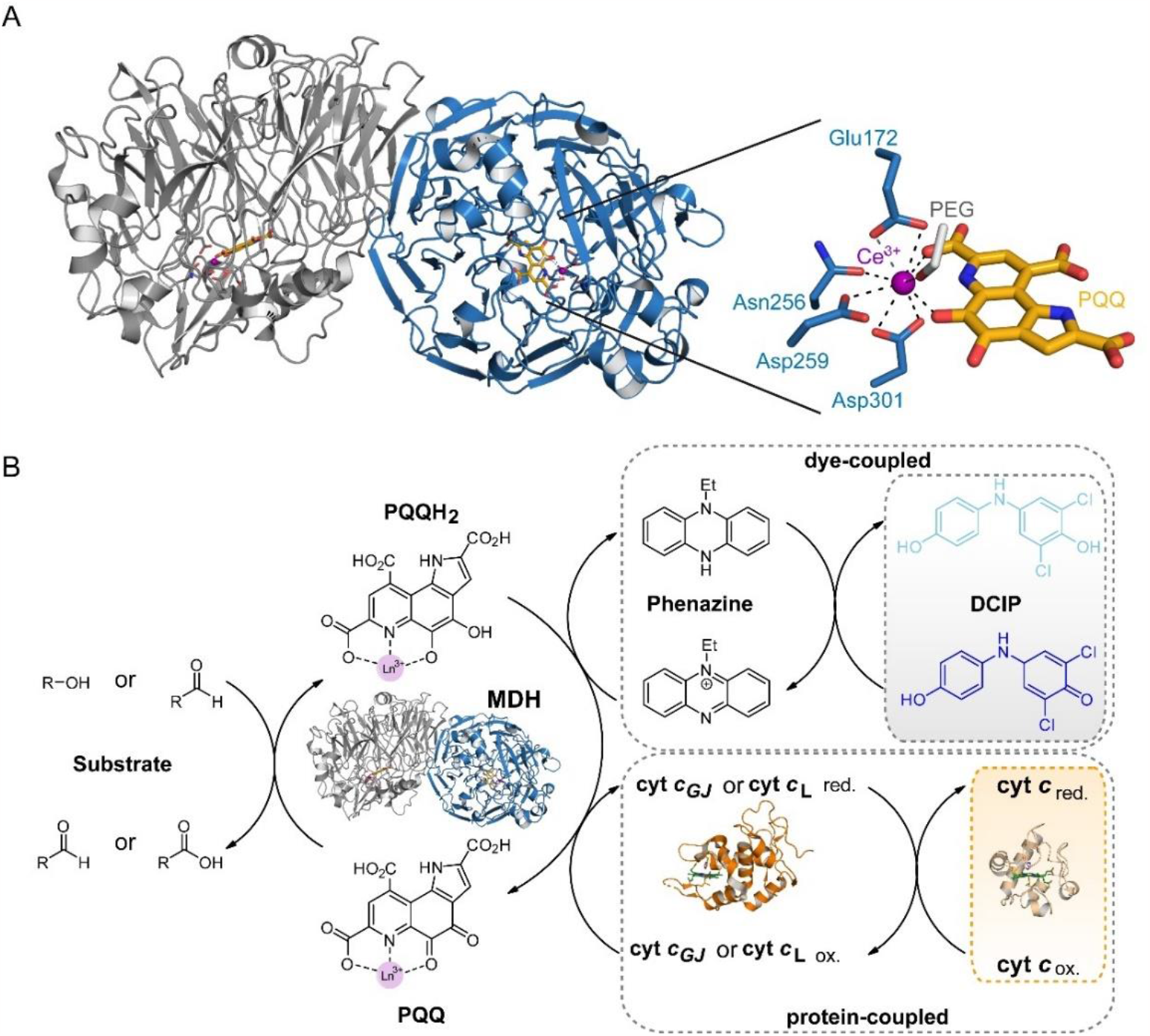
(A) Homodimeric structure of Ce-MDH from *Methylacidiphilum fuimariolicum* SolV (PDB: 4MAE)^20^ and zoom in its active center. The cofactors PQQ and Ce^3+^, as well as the coordinating amino acids are highlighted. The substrate binding coordination position of the Ce^3+^ is occupied by a polyethylene glycol (PEG) molecule from the crystallization buffer. (B) Schematic overview of the dye- and protein-coupled assay used in this study. Phenazine ethosulfate is reduced by PQQH_2_ in MDH, leading to the reduction of DCPIP and causing its discoloration. PQQH_2_ also reduces cyt *c*_*GJ*_ which in turn reduces equine heart cyt *c*. The change in absorbance for DCPIP and equine heart cyt *c* can be measured using UV/Vis at A_600_ and A_550_, respectively. The crystal structures of cytochrome *c* XoxG from *Methylorubrum extorquens* AM1 (PDB: 6ONQ) and equine heart cyt *c* (PDB: 1HRC) were used to illustrate this scheme. PQQ, pyrroloquinoline quinone. PQQH_2_, pyrroloquinoline quinol. DCPIP, 2,6-dichlorophenolindophenol. Modified from Refs. ^21^ and ^30^.

The mechanistic details of methanol oxidation by XoxF-MDH remains up for debate but two mechanisms are widely discussed by “wet-lab” and computational research groups: the addition-elimination and hydride transfer mechanism.^21-26^ Pol *et al*. isolated the acidophilic methanotroph *Methylacidiphilum fumariolicum* SolV from a volcanic mudpot.^[27]^ This bacterium is strictly dependent on Ln and exclusively possesses XoxF-MDH.^20, 27^ To cultivate strain SolV in a laboratory set-up, the extreme conditions of its natural environment have to be provided, including a Ln source, high temperature, low pH and the supply of methane and carbon dioxide.^20, 28, 29^ Most, if not all methylotrophic bacteria contain two distinct periplasmic, c-type cytochromes known as c_L_ and c_H_. In the past, these cytochromes were designated based on their isoelectric points (p*I*), cyt c_L_ having the lower p*I* value and cyt c_H_ the higher value. Only the cyt *c*_L_ is able to interact with MDH.^31^ The physiological electron transfer of MxaFI-MDH from substrate to its immediate cyt *c*_L_ was extensively studied by Anthony and co-workers in *Methylorubrum extorquens* AM1.^32-34^ Fundamentally, oxidation of methanol results in a concomitant reduction of PQQ to PQQH_2_, which is step-wise re-oxidized after two single electron transfers to two separate molecules of *c*_L_ which is reduced as well. Finally, cyt *c*_L_ regeneration is achieved through an additional electron transfer to cyt c_H_.^31^ In case of SolV, cyt *c*_L_ is termed cyt *c*_*GJ*_ and consists of the XoxG cytochrome and a periplasmic binding protein XoxJ.^35^ The interaction or “docking” of cyt *c*_*GJ*_ to XoxF-MDH involves electrostatic interaction between lysine residues on XoxF-MDH and carboxyl residues on cyt *c*_*GJ*_. Disassembly of the reduced cyt *c*_*GJ*_ is required before XoxF-MDH can interact with the next molecule of cyt *c*_*GJ*_. Electrostatic interactions are also involved for the electron transfer from cyt *c*_*GJ*_ to cyt *c*_H_.^32^ The interaction between XoxF-MDH and cyt *c*_*GJ*_ is disturbed by high salt concentration which inhibits substrate oxidation by up to 50% at 150 mM NaCl, 100 mM K_2_SO_4_ or 25 mM phosphate.^32, 35^ Furthermore, electrochemical experiments revealed that cyt *c*_*GJ*_ itself exhibits temperature dependence, with a 20% increase in current going from 10 to 35 °C.^36^ Featherston *et al*. characterized the immediate cytochrome of XoxF-MDH, XoxG, from *Methylorubrum extorquens* AM1. By comparing the results of an artificial dye-coupled assay with a XoxG-based assay, they concluded that these assays measure distinct aspects of XoxF-MDH activity.^37^

For decades, the most commonly practiced method to determine the activity of MDHs is a dye-coupled assay using the dyes phenazine ethosulfate (PES) as primary and 2,6-dichlorophenolindophenol (DCPIP) as secondary artificial electron acceptor.^38, 39^ The reduction of DCPIP results in its discoloration which is measured at 600 nm by UV/Vis. Based on previous work by Anthony and co-workers, protein-coupled activity assays for MDH from strain SolV using its physiological cyt *c*_*GJ*_ partner were developed.^30, 35, 40^ Commercially available cyt *c* from equine or bovine heart is used as secondary electron acceptor.^35^ Again, the reduction of the secondary electron acceptor can be monitored to determine the enzyme activity (Fig. 1B). Kalimuthu *et al*. developed an electrochemical assay that allows the co-adsorption of Eu-MDH and cyt *c*_*GJ*_ onto an electrode which functions as the secondary electron acceptor. In this case, the enzyme activity is tracked by measuring the current.^36^

Previous studies mainly used the dye-coupled assay to determine the enzyme activity of XoxF-MDH because the reagents are commercially available and low-cost. Usually a pH of 9, additives and activators like ammonia, glycine ethyl ester, potassium cyanide and an excess of Lns are included in the assay mixture to optimize the conditions for the assays (these additives are not necessary for MDH from *Methylacidiphilum fumariolicum* SolV).^30^ These experimental conditions rarely reflect the physiological conditions inside the cell and are truly artificial. Furthermore, the source of the endogenous substrate that causes background reaction is still unknown.^40^ Although, Featherston *et al*. compared the enzyme activity of La-, Ce- and Pr-MDH from *Methylorubrum extorquens* AM1 with the dye-coupled and protein-coupled assay, the differences in buffer, pH and activator do not allow for a direct comparison.^37^

Herein, we report the cultivation of *Methylacidiphilum fumariolicum* SolV with nine different Lns and the isolation and purification of the respective Ln-MDHs and their physiological electron acceptor cyt *c*_*GJ*_. Using the dye- and protein-coupled assays under the same assay conditions, we evaluated the enzyme activity of the different Ln-MDHs and observed varying trends among them, depending on the assay used.

## Results and Discussion

The growth rate of *Methylacidiphilum fumariolicum* SolV is highly dependent on supplemented Ln and its concentration in the growth medium.^20, 28^ Compared to the growth of SolV with early Ln La-Nd, the growth rate of SolV with late Ln Gd is less than half.^20^ When given a mixture of equimolar amount of all Ln and two actinides (Am, Cm), SolV preferably takes up the early Ln, showing the highest (80%) depletion from the medium for La.^28^ The depletion of Gd is barely 20% which is less than Am and Cm with a depletion of around 45% each.^28^ Based on the availability, XoxF-MDH is capable to incorporate a variety of Ln in its active site to obtain the respective Ln-MDH. We conducted nine separate cultivations of SolV at 55 °C and pH 2.7 with nine different Ln (La, Ce, Pr, Nd, Sm, Eu, Gd, Tb and Lu) in a self-build customized 3.5 L bioreactor, following a previously reported protocol.^29^ We observed exponential growth for seven out of nine cultivations (Fig. S1) and stopped the cultivation of SolV with Tb and Lu after 10 days as only linear growth was observed with these elements. Nonetheless, cells of all cultivations were collected for isolation of XoxF-MDHs and cyt *c*_*GJ*_. MDH makes up a high proportion of SolV’s biomass and can be isolated without an affinity tag as previously shown.^20^ The quantity of cyt *c*_*GJ*_ obtained after purification is low and makes native purification laborious. The heterologous expression of cyt *c*_*GJ*_ is desirable, but due to a so far unknown modification of or near its heme complex, native purification is, to the best of our knowledge, the only way to obtain cyt *c*_*GJ*_.^35^ Native protein purification of XoxF-MDHs and cyt *c*_*GJ*_ were performed by ion exchange chromatography and cyt *c*_*GJ*_ was additionally purified by size exclusion chromatography (SEC) (Fig. S2). The addition of 1 mM MeOH to all buffer solutions is imperative to ensure stability and activity of Ln-MDHs along the purification process and for long-term storage.^20^ To avoid secondary interactions of cyt *c*_*GJ*_ with the silica matrix, NaCl is added to the buffer for SEC. However, since the interaction of MDH and cyt *c*_*GJ*_ is mainly electrostatically and thus negatively affected by high salt concentrations, NaCl needs to be removed after SEC, which was done using centrifugal filter devices.^35^ Ln-MDHs and cyt *c*_*GJ*_ were analyzed by SDS-PAGE (Fig. S3) and the metal content of all Ln-MDHs was measured by Inductively Coupled Plasma Mass Spectrometer (ICP-MS).

ICP-MS measurement does not provide information about the metalation of the active site but rather the metal content of the sample. This can pose a challenge when XoxF-MDH co-purifies with other Ln binding proteins such as LanM rendering the read-out ineffective, as noted by Featherston *et al*.^37^ Strain SolV does not encode LanM in its genome and the use of time-resolved laser-induced fluorescence spectroscopy (TRLFS) enables the differentiation between Eu in the active site and in solution. TRLFS confirms the previously determined metal content value of Eu-MDH obtained by ICP-MS analysis.^41^ Previous studies found that XoxF-MDH from strain SolV is metallated on average with 60–70% of the respective Ln ions after purification.^20, 21, 41^ Our results show that the metal content of Ln-MDH is higher in the case of early Lns (La-Nd), ranging from 42–48%. However, there is a notable decrease in metalation across the Ln series, with Tb-MDH having 11% metal content. La-XoxF1 from *Methylorubrum extorquens* AM1 with a metal content of 39% displayed only half the specific activity compared to its previous studies.^42^ As the metal content is highly variable depending on the batch and metal, we recommend to determine the metal content of Ln-MDHs, before conducting any experiment to ensure accurate and reliable data. Despite their similar ionic radii, these small differences influence the incorporation efficiency and retention of the Ln in the active site. MDH obtained from strain SolV grown with Lu did not contain detectable amounts of Lu and was omitted for further experiments. PQQ is the second indispensable cofactor in the active site of Ln-MDH. The occupancy of the cofactor was determined by measuring its absorption maximum at 355 nm.^21, 22^ The cofactor was present in all samples but whether the loading is 100% cannot be taken from this method. Fully accounting for PQQ and/or metal contents continues to pose a challenge. Despite ongoing efforts, a comprehensive understanding of the exact mechanisms and factors influencing the presence and quantity of PQQ and/or metals remains elusive.

With eight Ln-MDHs and cyt *c*_*GJ*_ in hand, we moved on to determine the methanol oxidation activity. For decades, an artificial dye-coupled assay that utilizes redox-active dyes has been widely employed to assess the enzyme activity of MDHs derived from a vast variety of microorganisms.^1, 21, 43^ In addition to the non-specificity of the dye-coupled assay to MDH, the commercially available and low-cost reagents favor the usage of this assay. However, this method does not mirror the mechanism *in vivo* and conclusions drawn from the dye-coupled assay should be considered carefully. PES/DCPIP and the alternative dye-coupled method with Wurster’s Blue are all photosensitive, can induce high background reactivity and should be performed vigilantly.^39^ Similar to previous work, the specific activity of the Ln-MDHs with early Lns is higher and increases towards Nd-MDH and then decline subsequently (Fig. 2A). In order to account for the varying metal content among samples, we adjusted the specific activities by assuming a 100% metal content within the active site. Due to the relatively low metal content of Tb (11.3%) in the MDH sample purified from cells grown on Tb and likely even lower metalation of the active site and activation of PQQ, the adjusted specific activity is likely to be artificially inflated, but was included for transparency. For the smaller Lns, several theoretical studies have shown that the activation of PQQ is insufficient, preventing effective oxidation.^22, 44^ The low metalation of the sample leads to a substantial increase in uncertainty and will not be further discussed. Earlier studies obtained and compared the activity of different XoxF-MDHs by titrating the desired Ln to partial-apo Eu-MDH or incubating apo-MDH with Ln and PQQ.^21, 22, 28, 43^ To the best of our knowledge, the isolation of various Ln-MDHs from a microorganism grown with its respective Ln has not been widely practiced so far. Singer *et al*. were able to receive XoxF-MDH containing the heavier, smaller Lns by expressing an apo-XoxF-MDH in *Escherichia coli* and incubation with the respective Ln and PQQ for 72 hours.^28^ The results of their dye-coupled activity assays show that Tb-MDH is less active than Gd-MDH (although it should be noted, that metalation was not investigated here).

**Figure 2.**
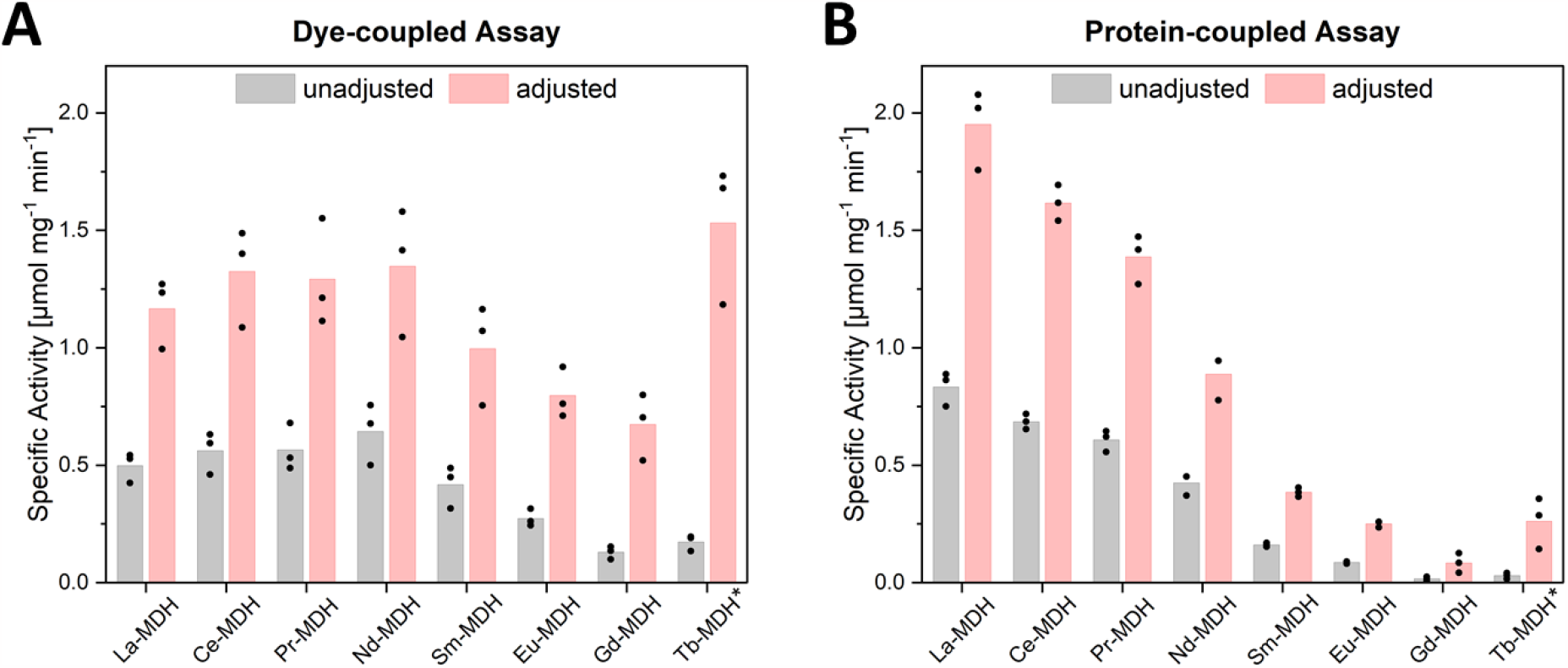
Results of the determination of enzyme activity using (A) the dye-coupled and (B) protein-coupled assay (outlined in Fig. 1B). Conditions dye-coupled assay: 100 nM XoxF-MDH, 1 mM PES, 100 μM DCPIP, 50 mM MeOH in 10 mM PIPES with 1 mM MeOH, pH 7.2, 45 °C. Conditions protein-coupled assay: 100 nM XoxF-MDH, 5 μM cyt *c*_*GJ*_, 50 μM cyt *c* from equine heart, 50 mM MeOH in 10 mM PIPES with 1 mM MeOH, pH 7.2, 45 °C. The adjusted values are calculated assuming 100% metal content in the active site. Each dot represents the result of one measurement. *The relatively low metal content of Tb-MDH (11.3%) likely inflates the adjusted value for the enzymatic activity.

Computational and experimental studies also discuss the effect of the Lewis acidity of the Lns on enzyme activity.^21-24^ Due to the Ln contraction, the Lewis acidity increases across the Ln series. PQQ requires a Lewis acid to activate its C5 quinone C–O bond for the subsequent proton abstraction step of the substrate. Higher Lewis acidity facilitates the rate-limiting breaking of the substrate C–H bond hence increasing substrate turnover and enzyme activity but obtaining XoxF-MDH with late Lns remains challenging.

A protein-coupled activity assay is another method to investigate kinetic parameters of MDHs. Anthony and co-workers developed a protein-coupled activity assay for MxaFI-MDH from *Methylorubrum extorquens* AM1.^40, 45^ The assay mixture is composed of MxaFI-MDH, its physiological partner cyt *c*_L_, cyt *c* from equine or bovine heart and MeOH as substrate. The physiological electron acceptor cyt *c*_L_ is used to re-oxidize MxaFI-MDH and the introduction of the secondary cytochrome *c* from equine or bovine heart is able to re-oxidze cyt *c*_L_ without interacting with MDH (Fig. 1B).^45^ Versantvoort *et al*. have shown with XoxF-MDH from SolV that the rate of reduction of the secondary cytochrome depends on the concentration of the physiological partner cytochrome which appeared to be linear between 0 and 1 μM for cyt *c*_*GJ*_.^35^ This method reflects the physiological mechanism more accurately and is easier to handle without photosensitive reagents. A limitation is the MDH-specificity of the real physiological cytochrome partners although some rare cases are reported where this was still possible, e.g. the cyt *c*_L_ of AM1 is able to interact with MDH from *Paracoccus denitrificans* and *Methylophilus methylotrophus*.^45^ Featherston *et al*. discussed the subtle differences of the immediate cytochromes MxaG and XoxG of MxaFI-MDH and XoxF-MDH, respectively, from *Methylorubrum extorquens* AM1.^37^ Both cytochromes are *c*-type cytochromes and carry the typical characteristics: covalent attachment of the heme *c* moiety to the protein via two thioether bonds and axial ligation of the Fe^3+^ by histidine and a second, in this case, a methionine residue.^46, 47^ The main distinctions are the loss of a helix in cyt *c* XoxG and the absence of a Ca^2+^ binding site in another helix. These differences result in more solvent-exposed heme that is proposed to be the source of the relatively low midpoint reduction value of the XoxG-type cytochromes.^37^ These subtle yet important structural changes are likely to result in different enzymatic activity of MDHs *in vivo* and cannot be properly reflected by an artificial dye-coupled assay. Thus, conclusions deduced from dye-coupled assays should be drawn carefully.

The highly laborious work to obtain adequate protein quantities required for protein-coupled activity assays is another hindrance, which should be considered. Before we conduct protein-coupled assay of XoxF-MDHs with cyt *c*_*GJ*_, we determined the optimal concentration of cyt *c*_*GJ*_ to obtain maximum substrate turnover. Among all tested Ln-MDHs, Nd-MDH demonstrated the highest enzyme activity based on the dye-coupled assay and was chosen as the proxy for this experiment. Nd-MDH was assayed with increasing amount of cyt *c*_*GJ*_ (0–30 μM) and Michaelis-Menten kinetics were obtained by best curve fitting (Fig. S4).

Based on these results, a concentration of cyt *c*_*GJ*_ above 100 μM is required to obtain v_max_, which is a prohibitively large amount of cyt *c*_*GJ*_ for a single assay. Therefore, we decided to choose a value at the end of the linear phase and proceeded with 5 μM cyt *c*_*GJ*_ for each assay. We evaluated the enzymatic activity of eight XoxF-MDHs with the protein-coupled assay and the results are shown in Fig. 2B. In order to exclude any day-to-day deviation, we used the same enzyme batch for both methods and conducted the experiments on the same day. In contrast to the dye-coupled assay, we observed a gradual decrease in enzymatic activity across the Ln series. Again, Gd-MDH displays the lowest enzyme activity, while La-MDH exhibits the highest enzyme activity. Regardless of method, Gd-MDH constantly remains the least active MDH. As discussed above, Lewis acidity can have a significant impact on substrate turnover through various mechanisms. Additional functions of the Lewis acid, like substrate orientation, cofactor redox cycling and substrate activation and their effects on the enzyme activity, have been discussed.^22, 24, 25, 48, 49^ We found that the specific enzyme activity of La-, Ce- and Pr-MDH is higher with the protein-coupled assay compared to their activity measured with the dye-coupled assay. In contrast, Nd-, Sm-, Eu- and Gd-MDH exhibit lower specific activity with the protein-coupled assay than with the dye-coupled assay. These results indicate that, depending on the assay, distinct trends can be observed among the XoxF-MDHs, implying that different aspects of the enzyme activity are being measured. Moreover, the protein-coupled assay revealed greater specific enzyme activity for La-, Ce- and Pr-MDH suggesting that these Ln may positively affect the efficiency in cyt *c*_*GJ*_ reduction. In comparison to the dye-coupled assay, protein-protein interaction steps are involved in the protein-coupled assay to transfer the electrons from XoxF-MDH to the final electron acceptor, but the positive effect of La, Ce and Pr on the interaction still exceed this challenge. These results indicate that La, Ce and Pr are not just suitable Lewis acids that catalyze the rate-limiting step for substrate turnover, but also facilitate and enhance the electron transfer between XoxF-MDH and its cyt *c*_*GJ*_, thereby increasing the overall functional efficiency. We cannot rule out that the use of 5 μM cyt *c*_*GJ*_ may not be optimal and might be sufficient to reach v_max_ for one XoxF-MDH but not for another such as Nd-MDH.

## Conclusions

To conclude, we have grown *Methylacidiphilum fumariolicum* SolV with nine different Lns and were able to purify eight different XoxF-MDHs, and their physiological electron acceptor cyt *c*_*GJ*_ and assessed the enzyme activity by performing dye- and protein-coupled activity assays. ICP-MS measurements revealed that XoxF-MDH contain varying amount of Ln in their active site and that this value decreases towards Tb. ICP-MS also showed that XoxF-MDH isolated from Lu-grown strain SolV was not able to incorporate any Lu. Next, we determined the enzyme activity using the dye-coupled assay with PES/DCPIP and compared the results with the protein-coupled assay using cyt *c*_*GJ*_ as electron acceptor. The data obtained from each method showed a discernible and distinct pattern, providing evidence that these methods are impacted by different aspects of the Ln-dependent enzyme activity. Both methods revealed that Gd-MDH is the least active among the eight XoxF-MDHs and that the enzymes containing larger Lns have higher activity, but different trends amongst the Ln series are observed based on the method used. At first glance, the increased activity of XoxF-MDH with early Lns align with the higher growth rates of strain SolV when cultivated with early Lns, suggesting that the growth of SolV is mainly linked to the activity of XoxF-MDH.^20, 28^ Wegner and co-workers discovered in *Beijerinckiaceae* bacterium RH AL1 that the addition of La or a Ln cocktail change the expression of nearly 41% of all genes in the genome and that different Lns affect different genes. Differentially expressed genes are associated with various biological processes of the Ln-dependent metabolism but also include secretion and uptake system, the flagellar and chemotactic machinery and other cellular functions.^50, 51^ Therefore, the influence of Lns extend beyond their catalytic role in methanol oxidation.

Using the dye-coupled assay, the enzyme activity increases towards Nd-MDH and declines afterwards. These results are in accordance with previous studies.^21, 22, 28, 43^ Using the protein-coupled assay, La-MDH displayed the highest enzyme activity which decreases progressively towards Gd-MDH. Although more steps are involved to transfer the electrons from XoxF-MDH to the final electron acceptor, we observed that La-, Ce- and Pr-MDH exhibit higher enzymatic activity in the protein-coupled assay than in the dye-coupled assay. This indicates that the enhancing effects of Lns are not limited to XoxF-MDH but also influence the electron transfer by its cyt *c*_*GJ*_. When adjusting the enzyme activity by their varying metal content, the trends became more pronounced, supporting our findings. This normalization allowed us to account for any potential discrepancies in the metal content and increases the accuracy and reliability of the results. To enhance reproducibility and comparability, we propose to incorporate the determination of metal content by ICP-MS into any kinetic experiments that involves metalloenzymes. This additional step will ensure that variations in metal content are accounted for, which will enable more meaningful comparisons between different experiments.

## Acknowledgments

The authors thank Christine Benning for ICP-MS measurements. We thank Dr. Arjan Pol for his input on SolV cultivation. SS, MTP and LJD acknowledge support from the Deutsche Forschungsgemeinschaft (DFG) projects 325871075 (SFB1309), 392552271 and SCHN1273-5.

## Notes

### Competing Interest Statement

The authors have declared no competing interest.

## References

(1) Nakagawa, T. et al. Catalytic Role of XoxF1 as La3+-Dependent Methanol Dehydrogenase in Methylobacterium extorquens Strain AM1. PLOS ONE 7, (11), e50480, DOI: 10.1371/journal.pone.0050480 (2012).

(2) Kato, S. et al. Isolation and Genomic Characterization of a Proteobacterial Methanotroph Requiring Lanthanides. Microbes Environ. 35, (1), DOI: 10.1264/jsme2.ME19128 (2020).

(3) Vekeman, B. et al. Genome Characteristics of Two Novel Type I Methanotrophs Enriched from North Sea Sediments Containing Exclusively a Lanthanide-Dependent XoxF5-Type Methanol Dehydrogenase. Microb. Ecol. 72, (3), 503–509, DOI: 10.1007/s00248-016-0808-7 (2016).

(4) Taubert, M. et al. XoxF encoding an alternative methanol dehydrogenase is widespread in coastal marine environments. Environ. Microbiol. 17, (10), 3937–3948, DOI: 10.1111/1462-2920.12896 (2015).

(5) Lv, H. et al. Isolation and genomic characterization of Novimethylophilus kurashikiensis gen. nov. sp. nov., a new lanthanide-dependent methylotrophic species of Methylophilaceae. Environ. Microbiol. 20, (3), 1204–1223, DOI: 10.1111/1462-2920.14062 (2018).

(6) Roszczenko-Jasińska, P. et al. Occurrence of XoxF-type methanol dehydrogenases in bacteria inhabiting light lanthanide-rich shale rock. FEMS Microbiol. Ecol. 97, (2), DOI: 10.1093/femsec/fiaa259 (2020).

(7) Hou, S. et al. Complete genome sequence of the extremely acidophilic methanotroph isolate V4, Methylacidiphilum infernorum, a representative of the bacterial phylum Verrucomicrobia. Biol. Direct 3, (1), 26, DOI: 10.1186/1745-6150-3-26 (2008).

(8) Chistoserdova, L. Modularity of methylotrophy, revisited. Environ. Microbiol. 13, (10), 2603–2622, DOI: 10.1111/j.1462-2920.2011.02464.x (2011).

(9) In Yeub, H. et al. Biocatalytic Conversion of Methane to Methanol as a Key Step for Development of Methane-Based Biorefineries. J. Microbiol. Biotechnol. 24, (12), 1597–1605, DOI: 10.4014/jmb.1407.07070 (2014).

(10) Hakemian, A. S. & Rosenzweig, A. C. The Biochemistry of Methane Oxidation. Annu. Rev. Biochem. 76, (1), 223–241, DOI: 10.1146/annurev.biochem.76.061505.175355 (2007).

(11) Kang, T. J. & Lee, E. Y. Metabolic versatility of microbial methane oxidation for biocatalytic methane conversion. J. Ind. Eng. Chem. 35, 8–13, DOI: 10.1016/j.jiec.2016.01.017 (2016).

(12) Schmitz, R. A. et al. Neodymium as Metal Cofactor for Biological Methanol Oxidation: Structure and Kinetics of an XoxF1-Type Methanol Dehydrogenase. mBio 12, (5), e01708–01721, DOI: doi:10.1128/mBio.01708-21 (2021).

(13) Chistoserdova, L. & Kalyuzhnaya, M. G. Current Trends in Methylotrophy. Trends Microbiol. 26, (8), 703–714, DOI: 10.1016/j.tim.2018.01.011 (2018).

(14) Mackenzie, C. et al. The home stretch, a first analysis of the nearly completed genome of Rhodobacter sphaeroides 2.4.1. Photosyn. Res. 70, (1), 19–41, DOI: 10.1023/A:1013831823701 (2001).

(15) Bosch, G. et al. Insights into the physiology of Methylotenera mobilis as revealed by metagenome-based shotgun proteomic analysis. Microbiology 155, (4), 1103–1110, DOI: 10.1099/mic.0.024968-0 (2009).

(16) Vu, H. N. et al. Lanthanide-dependent regulation of methanol oxidation systems in Methylobacterium extorquens AM1 and their contribution to methanol growth. J. Bacteriol. 198, (8), 1250–1259, DOI: 10.1128/jb.00937-15 (2016).

(17) Chu, F. & Lidstrom, M. E. XoxF Acts as the Predominant Methanol Dehydrogenase in the Type I Methanotroph Methylomicrobium buryatense. J. Bacteriol. 198, (8), 1317–1325, DOI: doi:10.1128/jb.00959-15 (2016).

(18) Gu, W. et al. Uptake and effect of rare earth elements on gene expression in Methylosinus trichosporium OB3b. FEMS Microbiol. Lett. 363, (13), DOI: 10.1093/femsle/fnw129 (2016).

(19) Masuda, S. et al. Lanthanide-Dependent Regulation of Methylotrophy in Methylobacterium aquaticum Strain 22A. mSphere 3, (1), 10.1128/msphere.00462-00417, DOI: doi:10.1128/msphere.00462-17 (2018).

(20) Pol, A. et al. Rare earth metals are essential for methanotrophic life in volcanic mudpots. Environ. Microbiol. 16, (1), 255–264, DOI: 10.1111/1462-2920.12249 (2014).

(21) Jahn, B. et al. Similar but Not the Same: First Kinetic and Structural Analyses of a Methanol Dehydrogenase Containing a Europium Ion in the Active Site. ChemBioChem 19, (11), 1147–1153, DOI: 10.1002/cbic.201800130 (2018).

(22) Lumpe, H. et al. Impact of the lanthanide contraction on the activity of a lanthanide-dependent methanol dehydrogenase – a kinetic and DFT study. Dalton Trans. 47, (31), 10463–10472, DOI: 10.1039/C8DT01238E (2018).

(23) Prejanò, M. et al. How Lanthanide Ions Affect the Addition–Elimination Step of Methanol Dehydrogenases. Chem. Eur. J. 26, (49), 11334–11339, DOI: 10.1002/chem.202001855 (2020).

(24) Prejanò, M. et al. How Can Methanol Dehydrogenase from Methylacidiphilum fumariolicum Work with the Alien CeIII Ion in the Active Center? A Theoretical Study. Chem. Eur. J. 23, (36), 8652–8657, DOI: 10.1002/chem.201700381 (2017).

(25) Leopoldini, M. et al. The Preferred Reaction Path for the Oxidation of Methanol by PQQ-Containing Methanol Dehydrogenase: Addition–Elimination versus Hydride-Transfer Mechanism. Chem. Eur. J. 13, (7), 2109–2117, DOI: 10.1002/chem.200601123 (2007).

(26) Anthony, C. The quinoprotein dehydrogenases for methanol and glucose. Arch. Biochem. Biophys. 428, (1), 2–9, DOI: 10.1016/j.abb.2004.03.038 (2004).

(27) Pol, A. et al. Methanotrophy below pH 1 by a new Verrucomicrobia species. Nature 450, (7171), 874–878, DOI: 10.1038/nature06222 (2007).

(28) Singer, H et al. Minor Actinides Can Replace Essential Lanthanides in Bacterial Life. Angew. Chem. Int. Ed. 62, e202303669, DOI: 10.1002/anie.202303669 (2023).

(29) Singer, H. et al. Learning from nature: recovery of rare earth elements by the extremophilic bacterium Methylacidiphilum fumariolicum. Chem. Comm., DOI: 10.1039/D3CC01341C (2023).

(30) Gutenthaler, S. M. et al./person-group>. in Methods in Enzymol. Vol. 650 (ed Cotruvo, J. A.), 57–79 (Academic Press 2021).

(31) Anthony, C. The c-type cytochromes of methylotrophic bacteria. Biochim. Biophys. Acta Bioenerg. 1099, (1), 1–15, DOI: 10.1016/0005-2728(92)90181-Z (1992).

(32) Cox, J. M. et al. The interaction of methanol dehydrogenase and its electron acceptor, cytochrome cL in methylotrophic bacteria. Biochim. Biophys. Acta Proteins Proteom. 1119, (1), 97–106, DOI: 10.1016/0167-4838(92)90240-E (1992).

(33) Anthony, C. Methanol Dehydrogenase, a PQQ-Containing Quinoprotein Dehydrogenase in Subcellular Biochemistry Vol. 35 (eds Holzenburg, A., Scrutton, N.S.), 73–117 (Springer US, 2000).

(34) Afolabi, P. R. et al. Site-Directed Mutagenesis and X-ray Crystallography of the PQQ-Containing Quinoprotein Methanol Dehydrogenase and Its Electron Acceptor, Cytochrome cL. Biochemistry 40, (33), 9799–9809, DOI: 10.1021/bi002932l (2001).

(35) Versantvoort, W. et al. Characterization of a novel cytochrome cGJ as the electron acceptor of XoxF-MDH in the thermoacidophilic methanotroph Methylacidiphilum fumariolicum SolV. Biochim. Biophys. Acta Proteins Proteom. 1867, (6), 595–603, DOI: 10.1016/j.bbapap.2019.04.001 (2019).

(36) Kalimuthu, P. et al. Electrocatalysis of a Europium-Dependent Bacterial Methanol Dehydrogenase with Its Physiological Electron-Acceptor Cytochrome cGJ. Chem. Eur. J. 25, (37), 8760–8768, DOI: 10.1002/chem.201900525 (2019).

(37) Featherston, E. R. et al. Biochemical and Structural Characterization of XoxG and XoxJ and Their Roles in Lanthanide-Dependent Methanol Dehydrogenase Activity. ChemBioChem 20, (18), 2360–2372, DOI: 10.1002/cbic.201900184 (2019).

(38) Ghosh, R. & Quayle, J. R. Phenazine ethosulfate as a preferred electron acceptor to phenazine methosulfate in dye-linked enzyme assays. Anal. Biochem. 99, (1), 112–117, DOI: 10.1016/0003-2697(79)90050-2 (1979).

(39) Jahn, B. et al. Understanding the chemistry of the artificial electron acceptors PES, PMS, DCPIP and Wurster’s Blue in methanol dehydrogenase assays. J. Biol. Inorg. Chem. 25, 199–212, DOI: 10.1007/s00775-020-01752-9 (2020).

(40) Day, D. J. & Anthony, C. Methanol dehydrogenase from Methylobacterium extorquens AM1. Meth. Enzymol. 188; 210–216, DOI: 10.1016/0076-6879(90)88035-9 (1990)

(41) Danaf, N. A. et al. Studies of pyrroloquinoline quinone species in solution and in lanthanide-dependent methanol dehydrogenases. PCCP 24, (25), 15397–15405, DOI: 10.1039/D2CP00311B (2022).

(42) Good, N. M. et al. Lanthanide-dependent alcohol dehydrogenases require an essential aspartate residue for metal coordination and enzymatic function. J. Biol. Chem 295, (24), 8272–8284, DOI: 10.1074/jbc.RA120.013227 (2020).

(43) Wehrmann, M. et al. Functional Role of Lanthanides in Enzymatic Activity and Transcriptional Regulation of Pyrroloquinoline Quinone-Dependent Alcohol Dehydrogenases in Pseudomonas putida KT2440. mBio 8, (3), e00570–00517, DOI: doi:10.1128/mBio.00570-17 (2017).

(44) Tsushima, S. Lanthanide-induced conformational change of methanol dehydrogenase involving coordination change of cofactor pyrroloquinoline quinone. PCCP 21, (39), 21979–21983, DOI: 10.1039/C9CP03953H (2019).

(45) Beardmore-Gray, M. et al. The methanol: cytochrome c oxidoreductase activity of methylotrophs. Microbiology 129, (4), 923–933, DOI: 10.1099/00221287-129-4-923 (1983).

(46) Bertini, I. et al. Cytochrome c: Occurrence and Functions. Chem. Rev. 106, (1), 90–115, DOI: 10.1021/cr050241v (2006).

(47) Liu, J. et al. Metalloproteins Containing Cytochrome, Iron–Sulfur, or Copper Redox Centers. Chem. Rev. 114, (8), 4366–4469, DOI: 10.1021/cr400479b (2014).

(48) Anthony, C. & Williams, P. The structure and mechanism of methanol dehydrogenase. Biochim. Biophys. Acta Proteins Proteom. 1647, (1), 18–23, DOI: 10.1016/S1570-9639(03)00042-6 (2003).

(49) Bogart, J. A. et al. DFT Study of the Active Site of the XoxF-Type Natural, Cerium-Dependent Methanol Dehydrogenase Enzyme. Chem. Eur. J. 21, (4), 1743–1748, DOI: 10.1002/chem.201405159 (2015).

(50) Wegner, C.-E. et al. Extracellular and Intracellular Lanthanide Accumulation in the Methylotrophic Beijerinckiaceae Bacterium RH AL1. Appl. Environ. Microbiol. 87, (13), e03144–03120, DOI: doi:10.1128/AEM.03144-20 (2021).

(51) Linda, G. et al. Different lanthanide elements induce strong gene expression changes in a lanthanide-accumulating methylotroph. bioRxiv, DOI: 10.1101/2023.03.06.530795 (2023).

