## Supporting Information for "Assessing Lanthanide-Dependent Methanol Dehydrogenase Activity: The Assay Matters"

### for

[1]Department of Chemistry, Ludwig-Maximilians-Universität München, Munich, Germany, [2]Chair of Analytical Chemistry, Technical University of Munich, Munich, Germany, [3]Department of Microbiology, Radboud Institute for Biological and Environmental Sciences, Faculty of Science, Radboud University, Nijmegen, The Netherlands, [4]Chair of Bioinorganic Chemistry, Heinrich-Heine-Universität Düsseldorf, Germany

Supporting figures, supporting table and experimental methods.

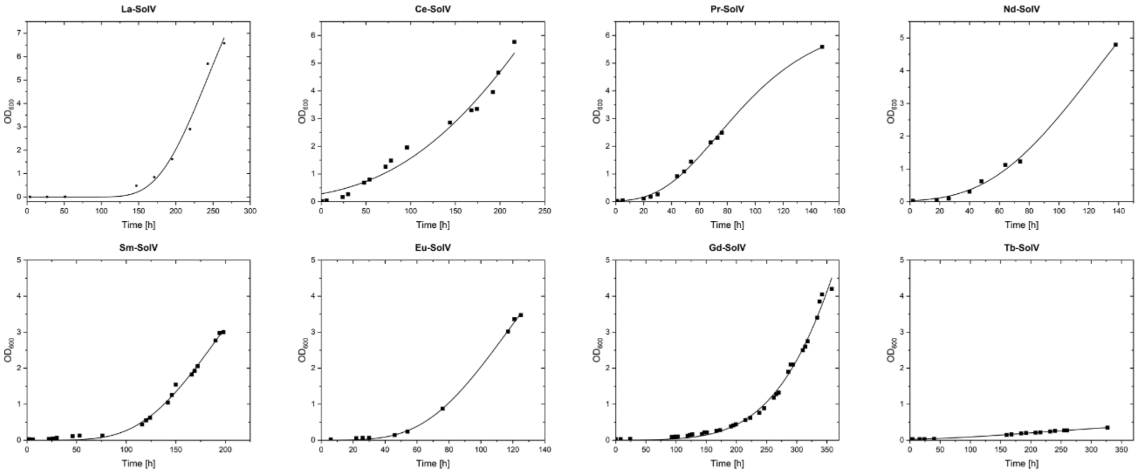


**Figure S1** Growth curve of SolV cultivated with different Lns (2 µM) at 55 °C. For composition of medium see Table S1. Protocol was followed as previously reported.[1]

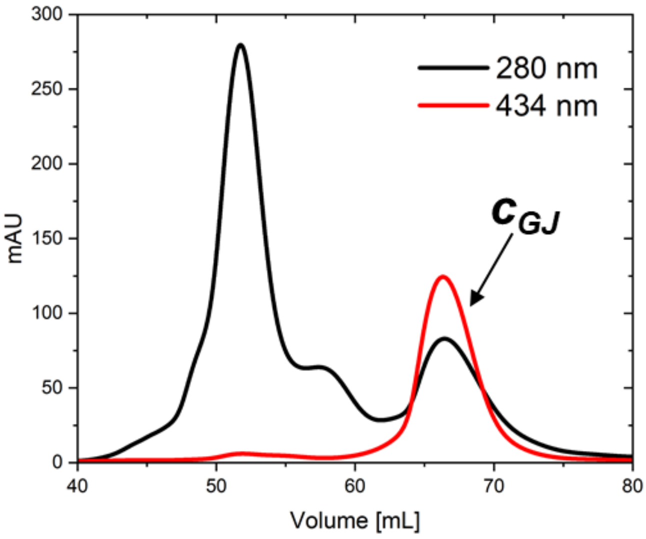


**Figure S2** Chromatogram of cyt $c_{GJ}$ purification by size exclusion chromatography (SEC). Conditions: 10 mM PIPES with 0.2 M NaCl, pH 7.2 at 4 °C.

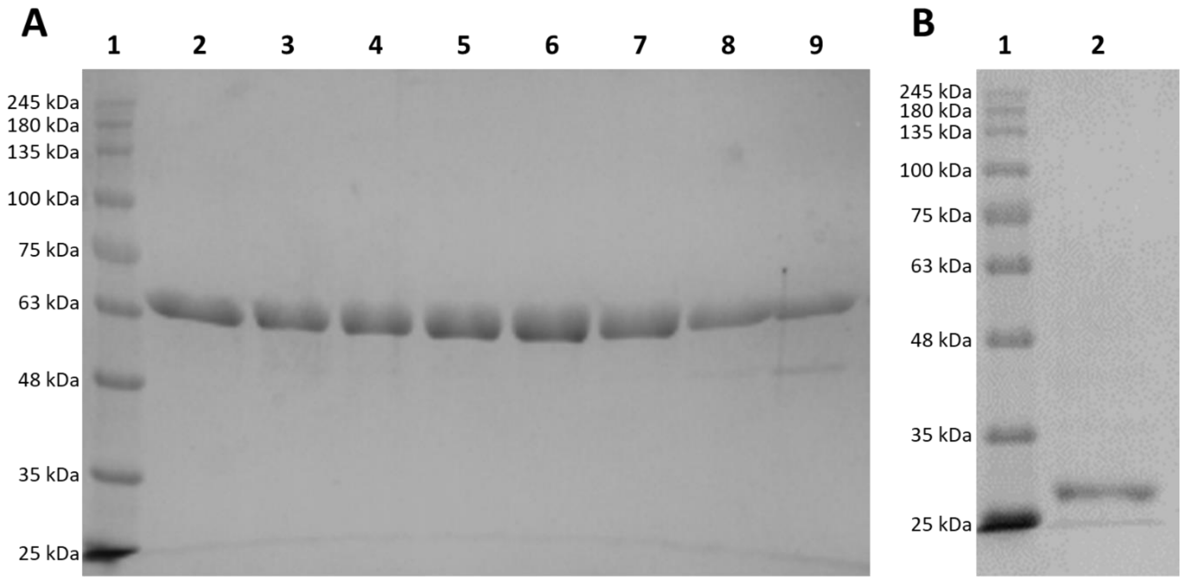


**Figure S3** SDS-PAGE analysis (12% w/v acrylamid) of (A) purified XoxF-MDHs (1: marker, 2: La-MDH, 3: Ce-MDH, 4: Pr-MDH, 5: Nd-MDH, 6: Sm-MDH, 7: Eu-MDH, 8: Gd-MDH, 9: Tb-MDH) and (B) purified cyt $c_{GJ}$ after size exclusion chromatography (1: marker, 2: cyt $c_{GJ}$). BlueEye prestained protein ladder (Jena Bioscience) was used as the marker.

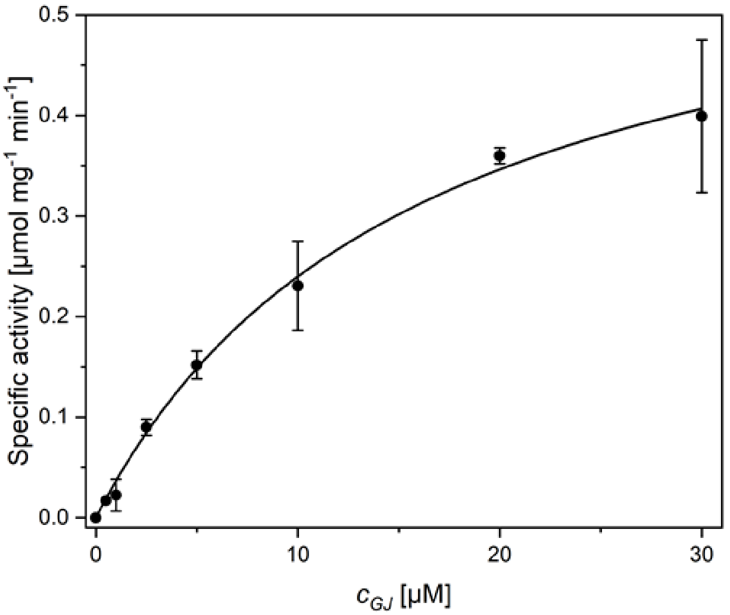


**Figure S4** Specific activity of Nd-MDH with increasing concentration of cyt $c_{GJ}$. Conditions: 100 nM Nd-MDH, 0–30 µM cyt $c_{GJ}$, 50 µM cytochrome *c* from equine heart, 50 mM MeOH in 10 mM PIPES with 1 mM MeOH, pH 7.2, 45 °C. Each dot represents the average of three replicates. $K_M$= 16.0 ± 1.9 µM, $v_{max}$= 0.62 ± 0.04.

**Table S1** Medium composition for the SolV growth medium.

| Solution | Composition |
|---|---|
| Minimal medium (10x) | 2 mM $MgCl_2 \cdot 6\ H_2O$, 10 mM $Na_2SO_4$, 20 mM $K_2SO_4$, 10 mM $NaH_2PO_4$, 2 mM $CaCl_2$<br>**Note:** All components except $CaCl_2$ were mixed and adjusted to pH 2.7 with 1 M $H_2SO_4$. $CaCl_2$ was autoclaved separately and added afterwards to prevent the precipitation of calcium phosphates. |
| Trace element (TE) solution | 200 mM $FeSO_4 \cdot 7\ H_2O$, 200 mM $MnCl_2 \cdot 4\ H_2O$, 300 mM $CuSO_4 \cdot 5\ H_2O$, 10 mM $NiCl_2 \cdot 6\ H_2O$, 10 mM $ZnSO_4 \cdot 7\ H_2O$, 10 mM $CoCl_2 \cdot 6\ H_2O$, 10 mM $Na_2MoO_4 \cdot 2\ H_2O$<br>**Note:** All components were dissolved one after the other in 1.5 % v/v $H_2SO_4$. Solution will occur blue immediately after preparation but turns green after a few days. |
| Growth medium for *M. fumariolicum* SolV | Prepared according to Pol *et al.*[2]<br>1x Minimal medium, 20 µL/L TE, 8 mM $NH_4^+$ for cultivation in bottles, for the bioreactor experiments the TE concentration was increased to 80 µL/L and 30 mM $NH_4^+$. All components were mixed and autoclaved.<br>**Note:** $NH_4^+$ available from a 2 M $(NH_4)_2SO_4$ solution in MilliQ water. |

### Methods

#### Bacterial culture

The cultivation of *Methylacidiphilium fumariolicum* SolV (La-Eu) was performed by using a modified protocol as previously reported.[3] For the composition of the growth medium see Table S1. SolV was grown with the desired lanthanide (La-Eu) in single-use polypropylene plastic cultivation flasks to an optical density at 600 nm ($OD_{600}$) of 0.5. Around 100–200 mL of strain SolV culture was used to inoculate the large-scale (3.5 L) bioreactor (non-commercial, self-build) to obtain a starting $OD_{600}$ of 0.05.[1] Throughout the cultivation, $CO_2$ (600 mL/min), $CH_4$ (750 mL/min) and air (1000 mL/min) was sparged through the medium. The temperature was kept at 55 °C and a stirring bar was used to ensure homogeneity and even distribution of gases.

The inoculants for Gd- and Tb-grown strain SolV were obtained by starvation of La-grown strain SolV through two cycles:

Minimal medium (100 mL) and La-grown strain SolV was added to a 1 L cultivation flask to a starting $OD_{600}$ of 0.05.

For incubation, a gas atmosphere of 85% air, 10% $CH_4$ and 5% $CO_2$ was provided. The flask was incubated in a shaker at 55 °C and 250 rpm for 4 days until an $OD_{600}$ of around 0.25 was reached (cycle 1). 100 mL of this starved La-grown strain SolV was used to repeat the same procedure once more until an $OD_{600}$ of 0.125 was reached (cycle 2). In the third round, 100 nM $GdCl_3$ or $TbCl_3$ was added to the starved strain SolV culture and further incubated until the desired $OD_{600}$ was reached and then used as inoculant for the bioreactor.

**Protein Purification**

*Methylacidiphilum fumariolicum* SolV cells were harvested by centrifugation at 8000 rpm for 10 min (Avanti JXN-26, Beckman Coulter). The cells were resuspended in 10 mM PIPES supplemented with 1 mM MeOH pH 7.2, and chemically lysed by commercially available BugBuster Protein Extraction Reagent (Merck, product code 70921). 10xBugBuster Protein Extraction Reagent was diluted to 1x with 10 mM PIPES and 1 mM MeOH pH 7.2. The reagent was added to frozen or thawed cell pellet (2 mL 1xBugBuster per 1 g resuspended cells), followed by the addition of 0.5–1.0 mg/mL lysozyme from chicken egg white (Sigma-Aldrich, CAS 12650-88-3), 0.2-0.5 mg/mL DNase I (PanReac AppliChem, product code A3778,0100) and incubated on a shaking platform for 30–45 min at room temperature. Afterwards, insoluble cell debris were removed by centrifugation (17 000 rpm, 20 min, 4 °C). After filtration of the supernatant with a filter paper (VWR, 5–13 µm particle retention), the sample was applied on a HiPrep™ SP Sepharose FF 16/10 cation exchange column (Cytiva, product code 28936544). The column was equilibrated with 10 mM PIPES, 1 mM MeOH pH 7.2 and bound proteins were eluted using 10 mM PIPES with 1 M NaCl and 1 mM MeOH pH 7.2. Cyt $c_{GJ}$ eluted at 2% (20 mM NaCl) and MDH at 25% (250 mM NaCl). Cyt $c_{GJ}$ was further concentrated using an Amicon® Ultra Centrifugal Unit (Merck, product code UFC901024) with a molecular weight cut-off of 10 kDa and applied on a HiLoad™ 16/600 Superdex™ 75 pg size exclusion column (Cytiva, product code 28989333). The column was equilibrated with 10 mM PIPES with 0.2 M NaCl pH 7.2 and cyt $c_{GJ}$ started to elute after 65 mL at a flowrate of 0.3 mL/min recording the UV/Vis absorption at 280 nm and its Soret peak of 434 nm. The NaCl concentration was reduced to less than 1 mM with an Amicon® Ultra Centrifugal Unit (Merck, UFC201024) with a molecular weight cut-off of 10 kDa and 10 mM PIPES pH 7.2.

For SDS-PAGE analysis, mPAGE® 4X LDS sample buffer (Merck, product code MPSB-10ML) containing 2% β-mercaptoethanol was added to samples and then heated at 70 °C for 5–7 min. The samples were loaded on a 12% SDS-PAGE gel (12% w/v acrylamide). The gel was stained with Coomassie Blue Stain for 1 h and subsequently treated with a destaining solution (10% (v/v) acetic acid and 20% (v/v) EtOH in ultrapure water). MDHs (63.5 kDa) and cyt $c_{GJ}$ (29.7 kDa) appear as dominant bands (Fig. S3).

**Activity Assays**

The activity assay with cyt $c_{GJ}$ and artificial electron acceptors are based on the protocols reported by Gutenthaler *et al.*[4] The enzymatic activity of MDH was assessed with native cyt $c_{GJ}$ through the reduction of equine heart cytochrome *c* (Sigma, CAS 9007-43-6). The reaction was monitored with an Epoch2 plate reader (formerly BioTek, now Agilent) at 45 °C through the increase of $A_{550}$. All experiments were conducted in 96-well-plates. Each well contained a total volume of 100 µL with

50 µM equine heart cytochrome *c*, 5 µM cyt $c_{GJ}$, 100 nM MDH and 50 mM MeOH in 10 mM PIPES with 1 mM MeOH pH 7.2. Everything but MDH were mixed together and incubated for 2 min at 45 °C before the reaction was initiated with the addition of MDH which was also incubated for 2 min at 45 °C. The extinction coefficient of equine heart cyt *c* was previously determined at 19.5 $mM^{-1}$ $cm^{-1}$ for 10 mM PIPES pH 7.2.[5] The specific activity was calculated using the slope of the initial 2 min after MDH addition. The specific activity of each experiment was adjusted according to the metal content of the MDH.

$$enzyme\ unit\ U\ [\mu mol\ min^{-1}] = \frac{initial\ rate\ of\ slope\ of\ measurement}{\varepsilon\ [cm^{-1}M^{-1}] \cdot pathlength\ of\ cell\ [cm]} \cdot 10^6 \cdot volume\ of\ assay\ [L]$$

$$specific\ activity\ [\mu mol\ min^{-1}\ mg^{-1}] = \frac{enzyme\ unit\ U\ [\mu mol\ min^{-1}]}{amount\ of\ enzyme\ [mg]}$$

$$specific\ activity_{adjusted}\ [\mu mol\ min^{-1}\ mg^{-1}] = \frac{enzyme\ unit\ U\ [\mu mol\ min^{-1}]}{amount\ of\ enzyme\ [mg] \cdot \frac{metal\ content\ [\%]}{100}}$$

The activity assay with the artificial electron acceptor DCPIP (2,6-Dichlorophenolindophenol sodium salt dihydrate, formerly Fluka, now Honeywell, CAS: 1266615-56-8) and PES (phenazine ethosulfate, Sigma-Aldrich, CAS 10510-77-7) was assessed through the reduction of DCPIP. The reaction was monitored with an Epoch2 plate reader (formerly BioTek, now Agilent) through the decrease of $A_{600}$. Each well contained a total volume of 100 µL with 1 mM PES, 100 µM DCPIP, 100 nM MDH, 50 mM MeOH in 10 mM PIPES with 1 mM MeOH pH 7.2. Everything but MDH were mixed and incubated for 2 min at 45 °C in the dark before the reaction was initiated with addition of MDH which was also incubated for 2 min at 45 °C. The extinction coefficient of DCPIP was determined at 19.8 $mM^{-1}$ $cm^{-1}$ for 10 mM PIPES pH 7.2. The specific activity was calculated using the slope of the initial 2 min after MDH addition. The specific activity of each experiment was adjusted according to the metal content of the MDH.

$$enzyme\ unit\ U\ [\mu mol\ min^{-1}] = \frac{-1 \cdot initial\ rate\ of\ slope\ of\ measurement}{\varepsilon\ [cm^{-1}M^{-1}] \cdot pathlength\ of\ cell\ [cm]} \cdot 10^6 \cdot volume\ of\ assay\ [L]$$

$$specific\ activity\ [\mu mol\ min^{-1}\ mg^{-1}] = \frac{enzyme\ unit\ U\ [\mu mol\ min^{-1}]}{amount\ of\ enzyme\ [mg]}$$

$$specific\ activity_{adjusted}\ [\mu mol\ min^{-1}\ mg^{-1}] = \frac{enzyme\ unit\ U\ [\mu mol\ min^{-1}]}{amount\ of\ enzyme\ [mg] \cdot \frac{metal\ content\ [\%]}{100}}$$

The Michaelis-Menten constant $K_M$ and the maximum turnover speed $v_{max}$ of the cyt $c_{GJ}$-based activity assay (0–30 µM) with 100 nM Nd-MDH were calculated with the Michaelis-Menten equation using the slope of the initial 2 min after initiation:

$$v_0 = \frac{v_{max}[\mathrm{S}]}{K_\mathrm{M} + [\mathrm{S}]}$$

With $v_0$ representing the initial velocity and [S] the substrate concentration. The specific activity of each experiment was adjusted according to the metal content of Nd-MDH (47.8%).

**Metal analysis by ICP-MS**

The Ln-content of each MDH was determined by addition of the samples to 3% nitric acid (Suprapur®, Supelco) and heating for 1 h at 90 °C before analysis using an Inductively Coupled Plasma Mass Spectrometer (Nexion 350D, Perkin Elmer). Protein concentration were determined spectrophotometrically at 280 nm using an extinction coefficient of 158 $cm^{-1}$ $mM^{-1}$.[2] The metal content of the Ln-MDHs used in these experiments was determined at 42.7% for La-MDH, 42.4% for Ce-MDH, 43.8% for Pr-MDH, 47.8% for Nd-MDH, 41.9% for Sm-MDH, 34.3% for Eu-MDH, 19.2% for Gd-MDH and 11.3% for Tb-MDH.

### References

(1) Singer, H. *et al.* Learning from nature: recovery of rare earth elements by the extremophilic bacterium Methylacidiphilum fumariolicum. *Chem. Comm.* **59**, 9066-9069, DOI: 10.1039/D3CC01341C (2023).
(2) Pol, A. *et al.* Rare earth metals are essential for methanotrophic life in volcanic mudpots. *Environ. Microbiol.* **16**, 255-264, DOI: 10.1111/1462-2920.12249 (2014).
(3) Singer, H. *et al.* Minor Actinides Can Replace Essential Lanthanides in Bacterial Life. *Angew. Chem. Int. Ed.* **62**, DOI: 10.1002/anie.202303669 (2023).
(4) Gutenthaler, S. M. *et al.* in *Methods in Enzymol.* Vol. 650 (ed Cotruvo, J. A.), 57-79 (Academic Press 2021).
(5) Versantvoort, W. *et al.* Characterization of a novel cytochrome $c_{GJ}$ as the electron acceptor of XoxF-MDH in the thermoacidophilic methanotroph Methylacidiphilum fumariolicum SolV. *Biochim. Biophys. Acta Proteins Proteom.* **1867**, 595-603, DOI: 10.1016/j.bbapap.2019.04.001 (2019).
